# Identification and mapping of *Rpi-blb4* in diploid wild potato species *Solanum bulbocastanum*

**DOI:** 10.1101/2023.05.10.540114

**Authors:** Jie Li, Amanpreet Kaur, Brian Harrower, Miles Armstrong, Daolong Dou, Xiaodan Wang, Ingo Hein

## Abstract

More than 170 years after causing the potato famine in Ireland, late blight is still considered one of the most devastating crop diseases. Commercial potato breeding efforts depend on natural sources of resistance to protect crops from the rapidly evolving late blight pathogen, *Phytophthora infestans*. We have identified and mapped a novel broad-spectrum disease resistance gene effective against *P. infestans* from the wild, diploid potato species *Solanum bulbocastanum*. Diagnostic resistance gene enrichment sequencing (dRenSeq) was used to confirm the uniqueness of the identified resistance. RenSeq and GenSeq-based mapping of the resistance, referred to as *Rpi-blb4*, alongside recombinant screening, positioned the locus responsible for the resistance to potato Chromosome 5. The interval spans approximately 2.3 MB and corresponds to the DM reference genome positions of 11.25 and 13.56 Mb.

## 1. Introduction

Potato crop losses caused by the late blight pathogen *Phytophthora infestans* led to the Irish famine in the 1840s which resulted in the death of over one million people and the emigration of over two million throughout the world. Over a century after the famine, *P. infestans* remains one of the biggest threats to the global potato industry [1]. At present, disease control measures are highly dependent on the use of chemicals. The estimated annual cost associated with fungicide sprays and yield losses exceeds nine billion Euros [2]. However, the development of fungicide-resistant isolates and the introduction of stricter fungicide regulatory measures make chemical-based blight management less sustainable [3-5].

Utilising natural host resistances that are available in wild potato species is considered an important and environmentally benign disease control strategy [6,7]. Advances in potato genomics and marker developments can further expedite the introgression of resistance (*R*) genes from the wild species into commercial potato cultivars [8]. To counter the emergence of virulent *P. infestans* populations that can overcome such deployed host resistances, the continuous identification of novel genes and resistance sources is important for sustainable crop production.

A substantial description, identification, mapping, and/or allele mining of late blight resistance (*Rpi*) genes has been reported from Solanum species to date. This includes, for example, *R1*-*R11* from *S. demissum* [9-12], *Rpi-mcd1* from *S. microdontum* [13], *Rpi-mcq1* from *S. mochiquense* [14], *Rpi-phu1* from *S. phureja* [15], *Rpi-vnt1*.*1, Rpi-vnt1*.*2*, and *Rpi-vnt1*.*3* from *S. venturi* [15-17], *Rpi-pur1* from *S. piurae* [17], *Rpi-Smira 1* from *S. tuberosum* cv. Sapro Mira [8], *Rpi-tar1* and *Rpi-tar1*.*3* from *S. tarijense* [18,19], *Rpi-ver1 from S. verrucosum* [20], and many more that have recently been summarised by Paluchowska et al. [2]. Cloned *Rpi* genes typically belong to the group of nucleotide-binding, leucine-rich-repeat (NLR) genes.

The Commonwealth Potato Collection (CPC), based at the James Hutton Institute, forms part of an international potato GenBank network of potatoes. The CPC contains approximately 1500 accessions of wild and cultivated potato species including accessions of *S. bulbocastanum*. During the annual rejuvenation efforts, the CPC is routinely screened for the presence of novel blight resistances. Like the aforementioned cloned *Rpi*-genes, the screening of the CPC focuses on the identification of novel NLRs [21,22]. We developed <u>R</u>esistance gene <u>en</u>richment and <u>seq</u>uencing (RenSeq) technology to specifically sequence NLRs [23]. This targeted enrichment technology, in combination with Generic-mapping enrichment sequencing (GenSeq) and diagnostic RenSeq (dRenSeq) has, for example, enabled the mapping of a novel resistance, *Rpi-ver1*, from *S. verrucosum* [20, 22, 24]. Further, using RenSeq in combination with long-read sequencing technologies, functional NLRs such as *Rpi-amr1, Rpi-amr3i*, and *Rpi-amr3*, effective against late blight have been isolated from the non-tuber bearing species *S. americanum* [25-27].

Here we describe the characterisation of a novel resistance from *S. bulbocastanum*, a tuber-bearing, diploid wild potato species originating from Mexico and Guatemala which is known for its durable late blight resistance [28]. So far, five resistance genes, some of which are closely related, have been isolated from *S. bulbocastanum*. These comprise *Rpi-blb1/RB* and the homolog *Rpi-bt1* on chromosome 8 [29-31], *Rpi-blb2* on chromosome 6 [28], and *Rpi-blb3* (an *R2* homolog) on chromosome 4 [32]. In this study, we describe a new resistance in the *S. bulbocastanum* accession 7650 which we refer to as *Rpi-blb4*. We combined dRenSeq to ascertain the uniqueness of the resistance in the accession 7650 with RenSeq and GenSeq-based bulked segregant analysis of a population derived from a cross between the resistant plant *S. bulbocastanum* 7650 with *S. michoacanum* accession 3847 to genetically map this new resistance to potato chromosome 5.

## 2. Material and Methods

### 2.1. Plant material

Accessions of *Solanum bulbocastanum* that are maintained as part of the Commonwealth Potato Collection (CPC), were screened for late blight resistance. The screening revealed that *S. bulbocastanum* accession 7650, hereafter referred to as BLB7650, is highly resistant to late blight. A population that segregates for the resistance from BLB7650 was generated by crossing this accession with *S. michoacanum* accession MCH3847, a naturally occurring hybrid between *S. bulbocastanum* and *S. pinnatisectum* [33]. To map the resistance, 100 F1 progeny clones were established and evaluated.

### 2.2. Late blight pathogenicity assessment

Leaves from BLB7650, MCH3847, and the BLB7650 x MCH3847 progeny were tested in independent replicates for blight resistance through detached-leaf assay according to the method described by Whisson et al. [34] using the *Phytophthora infestans* isolate 2009-7654A. To assess the broadness of the resistance in BLB7650, 15 additional diverse *Phytophthora infestans* isolates were used. Disease severity was recorded on the Malcolmson scale [35] 5-8 days post-inoculation (dpi), where 1 represents highly susceptible and 5 highly resistant, respectively.

### 2.3. RenSeq and GenSeq-based genotyping

Genomic DNA from the parents BLB7650 and MCH3847 was extracted alongside 23 resistant (scoring 4 to 5) and 24 susceptible (scoring 1 to 2) progeny clones which were pooled to form the resistant and susceptible bulks. RenSeq and GenSeq were carried out using the method described previously [20]. Genomic DNA (1.0 μg) isolated from the parents and F1 individuals that were combined to form the bulks was fragmented using the Ultrasonicator M220 (Covaris) with peak power adjusted to 50W and cycles/burst to 200. Fragmented DNA was subjected to end-repair and each parent and bulk was individually barcoded with Illumina adaptors (NEBNext Ultra DNA library prep kit for Illumina) and hybridized with biotinylated RNA baits [23]. Post-enrichment, the indexed resistant/susceptible parents and bulks were sequenced on an Illumina MiSeq v2 sequencing system [20].

For dRenSeq and RenSeq/GenSeq analyses, paired-end Illumina MiSeq reads (2×250bp) were adapter trimmed according to the method described [22]. The trimmed RenSeq reads were mapped using Bowtie 2 [36] to a reference of known NB-LRR genes [24] including 5’ and 3’ sequences flanking the coding DNA sequence (CDS). In diagnostic mode, bowtie2 settings were adjusted so that only reads identical to the reference sequences are mapped (--score-min L, -0.01, -0.01). After mapping, the percentage coverage of each known NB-LRR gene was calculated with the expectation that if an accession contains a known NB-LRR gene this would be reflected by 100% read coverage.

### 2.4. Bulk Segregant analysis (BSA) and SNP filtering

The reads obtained from Illumina sequencing of separately indexed resistant/susceptible bulks and parents were mapped to the reference genome (*S. tuberosum* group Phureja clone DM1-3 516 R44 DM_V4.03; commonly known as DM) using bowtie 2 at 3%, 5% and 7% mismatch rates (--score-min L,-0.18,-0.18, L,-0.3,-0.3 and L,-0.42,-0.42). Consequently, single nucleotide polymorphisms (SNPs) between the reads and the reference sequence were called. SNPs were identified using samtools mpileup and Varscan v2.3.7. SNPs were filtered using a custom awk script for those SNPs that were present in both the parents and bulks at frequencies consistent with being linked to the resistant trait [20]. Specifically, a SNP was defined as being linked if it was absent in the susceptible parent and present at 50.0 % in the resistant parent while being absent in the susceptible bulk and present at 50.0 % in the resistant bulk. Conversely, a linked SNP present at 100.0 % in the susceptible parent should be present at 50.0 % in the resistant parent, 100.0 % in the susceptible bulk, and 50.0 % in the resistant bulk. To allow a margin of error due to sequences derived from closely related alleles or paralogs, a 10.0 % deviation from these expected values was permitted. The minimum read depth was set to 50.0 reads to reduce sampling artifacts. SNPs found at frequencies consistent with linkage to the resistance were further filtered for those that coincide within the coding sequences of NB-LRRs [37].

### 2.5. KASP assay for genotyping

Competitive allele-specific PCR (KASP) was used for marker-based genotyping. KASP primers were designed from the informative RenSeq and GenSeq SNPs linked to the resistance by providing 50 bp flanking sequence at the 5’ and 3’ end of the SNP, respectively. KASP-By-Design primers and KASP V4.0 2X Master mix (P/N KBS-1016-002 LGC Genomics limited) were used for the assays.

The KASP assays were conducted on a StepOne Plus real-time PCR machine using a total of 8.11 μl reaction mix which contained 20-40ng DNA, KASP v4.0 Reagent (KBS-1016), and KBD assay mix (KBS-1013) in working concentrations. The PCR program and genotype determination were operated as described [20].

## 3. Results

A screening of the Commonwealth Potato Collection for late blight-resistant accessions of *Solanum bulbocastanum* identified *S. bulbocastanum* accession 7650 (BLB7650) as resistant against the *Phytophthora infestans* isolate 2009-7654A, which belongs to the genotype 13_A2. The accession BLB7650 showed further strong resistance against 14 diverse *P. infestans* isolates from the U.S.A, The Netherlands, and the U.K. which indicates a broad-spectrum effectiveness of the resistance. The average scores against these isolates ranged from 4.3 to 5. Only one isolate, NL09096 from the Netherlands, yielded a score of 2 which is consistent with virulence and thus indicates an ability to overcome the resistance in BLB7650 (Table S1).

### 3.1. A novel resistance in *Solanum bulbocastanum* accession 7650, *Rpi-blb4*, controls late blight disease

To ascertain if the resistance in BLB7650 could be explained by the presence of already characterised NB-LRRs, a dRenSeq analysis was performed using the approach described previously [24]. Illumina-based RenSeq reads from BLB7650 were mapped to the individual reference genes under highly stringent conditions. Except for *Rpi-blb3*, no sequence identical to known NB-LRRs was found (Fig. 1; Table S2). *Rpi-blb3* belongs to a larger NLR gene family comprising functional homologs such as *R2, R2-like, Rpi-abpt*, and *Rpi-mcd1* on Chromosome 4 [32]. However, it has been demonstrated previously that the blight isolates from the genotype 13_A2 can typically overcome this resistance [38]. In line with this, *Rpi-blb3* was also present in progeny plants that were part of the resistant and susceptible bulks but not in MCH3847 (Fig. 1; Table S2). Due to the presence of *Rpi-blb3* in susceptible progeny, we conclude that a novel gene explains the resistance in BLB7650, and we refer to this gene as *Rpi-blb4*.

**Fig. 1.**
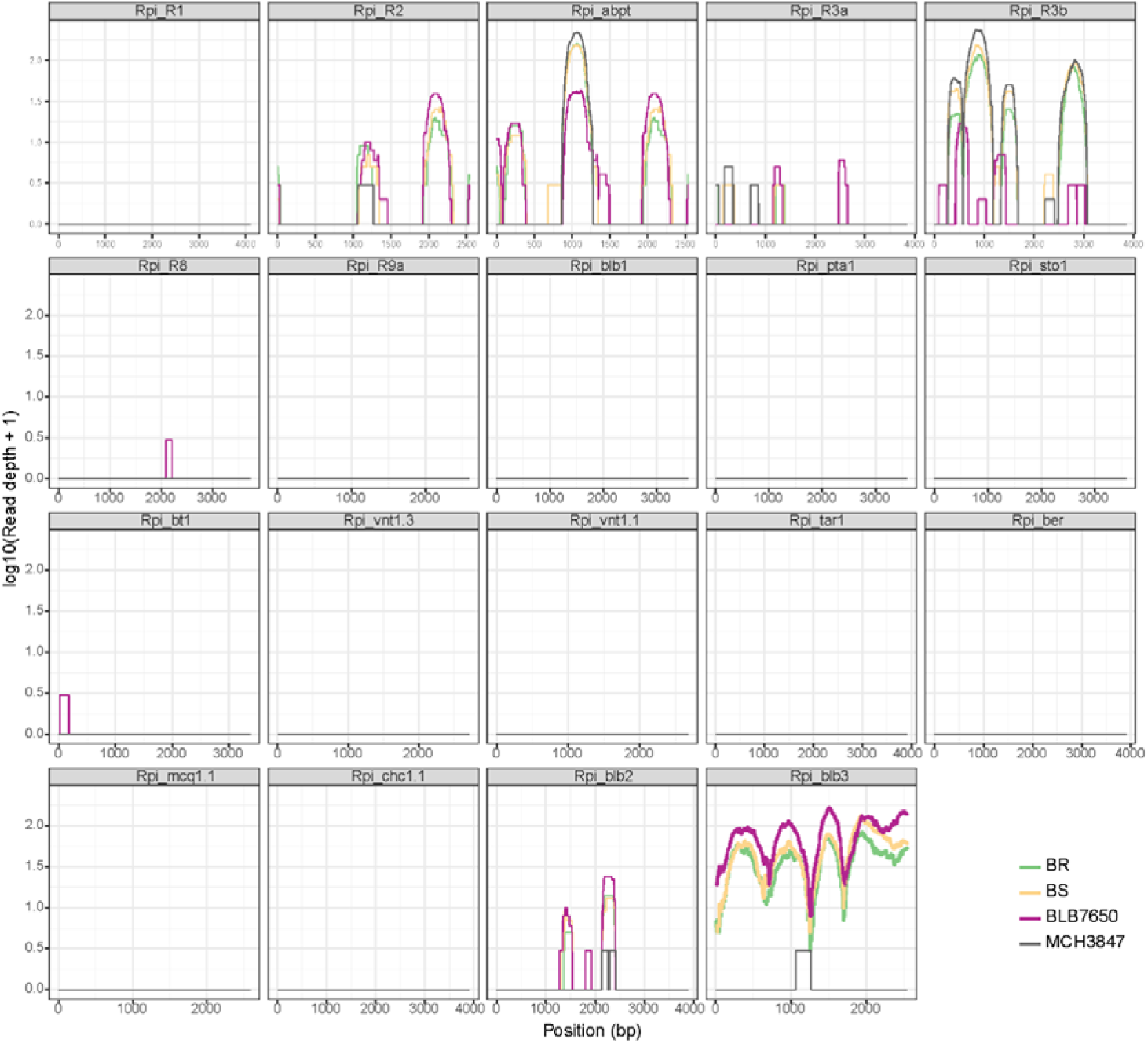
dRenSeq analysis of *Solanum bulbocastanum* accession 7650, *S. michoacanum* MCH3847, and resistant/susceptible progeny bulks. RenSeq-derived reads are mapped to the reference sequences of known NLR genes using stringent conditions. The x-axis of each box represents the whole sequence of the individual NLR from start to stop, while the coverage depth of each NB-LRR is shown on the y-axis on a log scale.

### 3.2. The resistance from BLB7650 segregates 1:1 in an F1 progeny derived from BLB7650 X MCH3847

After establishing that BLB7650 contains a novel resistance against *P. infestans* distinct from, for example, *Rpi-blb1/bt* and *Rpi-blb2* identified in *S. bulbocastanum* before, we established a population by crossing BLB7650 with MCH3847. In total, 100 seedlings of the F1 population were raised and assessed for late blight resistance using the *P. infestans* isolate 2009-7654A in independent replicates (Table S3). The average disease severity was calculated and plants scoring between 1 and <2.5 on the Malcolmson disease scale were designated as susceptible, progenies between 2.5 and <3.5 as intermediate, and plants scoring >3.5 as resistant (Table S3; Fig. 2). Out of 100 seedlings tested, 36 were susceptible, 19 were categorised as intermediate resistant/susceptible, and 45 as resistant. The approximate 1:1 ratio of resistant versus susceptible was confirmed through a Chi-Square test which yielded a Chi-Square value of 0.553856 with 1 degree of freedom and a *p*-value of 0.4567. The observed bimodal distribution of the contrasting phenotypes (Fig. 2) is consistent with the segregation of a single dominant resistance gene in this population, indicating that BLB7650 is most likely heterozygous for this resistance.

**Fig. 2.**
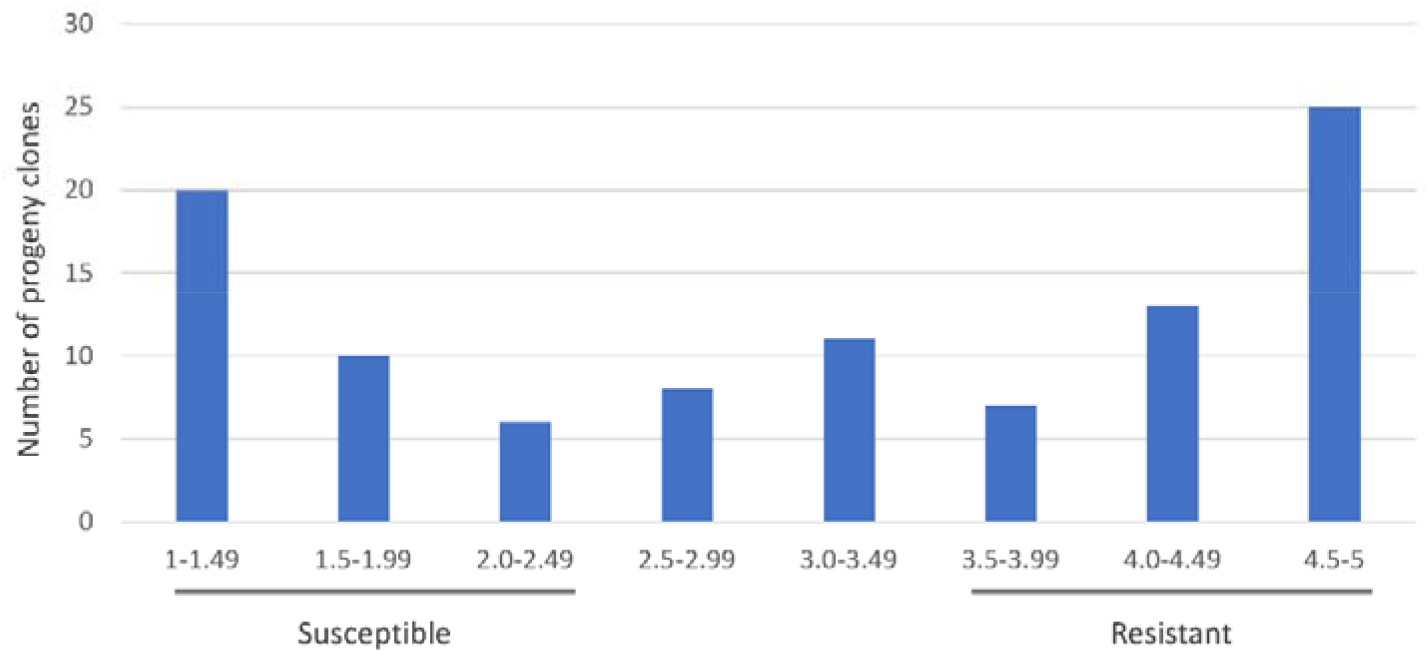
Late Blight score distribution of 100 F1 progeny clones derived from a cross between *S. bulbocastanum* accession 7650 and *S. michoacanum accession* MCH3847. Replicated detach leaf assays were used to assess the resistance of clones against the *P. infestans* isolate 2009-7654A. Scores between 1 and <2.5 are associated with susceptibility and scores >3.5 with resistance.

### 3.3. RenSeq and GenSeq analyses of the F1 progeny places *Rpi-blb4* on chromosome 5

Following the phenotypic assessment of the segregation population, bulks were compiled to reflect the most resistant and susceptible progeny clones. The bulked resistant sample comprised 23 clones all with a late blight score of over 4 and the bulked susceptible samples 24 clones with a score of equal or less than 2 (Table S3). Equimolar amounts of DNA from the selected resistant and susceptible progeny clones were combined to form the respective bulks and subjected to targeted enrichment RenSeq and GenSeq genotyping alongside the parents, BLB7650 and MCH3847.

SNPs that correspond in their frequency with the phenotypic evaluation that concluded a single dominant gene being responsible for the phenotype, were called at 3%, 5%, and 7% mismatch rates between the parents, the bulks, and the combination of the parents and bulks. At the 3% mismatch rate, 36 RenSeq and 10 GenSeq-derived SNPs were obtained in the coding sequences of the queried NLRs (RenSeq) or mainly single-copy genes (GenSeq). At a 5% mismatch rate, this increased to 58 and 57 SNPs respectively and yielded 75 and 66 SNPs at the 7% mismatch rate (Table 1).

**Table 1.** Single nucleotide polymorphism (SNP) analysis of reads obtained from RenSeq and GenSeq at different stringency conditions (3%, 5%, 7% mismatch rates)

| Mismatch<br>rates (%) | Technique | Parental<br>Filtered<br>SNPs | Parental<br>Filtered<br>SNPs in<br>NLRs | Bulk<br>Filtered<br>SNPs | Bulk<br>Filtered<br>SNPs in<br>NLRs | Parental<br>validated<br>filtered<br>SNPs | Parental<br>validated<br>filtered<br>SNPs in<br>NLRs |
| --- | --- | --- | --- | --- | --- | --- | --- |
| 3 | RenSeq | 1048 | 929 | 54 | 53 | 37 | 36 |
|  | GenSeq | 1169 | 1136 | 26 | 24 | 11 | 10 |
| 5 | RenSeq | 2884 | 2544 | 104 | 99 | 58 | 58 |
|  | GenSeq | 3041 | 2937 | 90 | 83 | 62 | 57 |
| 7 | RenSeq | 4475 | 3776 | 162 | 154 | 80 | 75 |
|  | GenSeq | 4279 | 4086 | 99 | 92 | 72 | 66 |

The genome of the *S. tuberosum* group Phureja clone DM 1-3 516 R44 (commonly known as DM) served as a reference for both the RenSeq and GenSeq analyses and was used to position the SNPs that are linked to the phenotype to the individual linkage groups. At the 3% mismatch rate, the 36 RenSeq-derived SNPs were derived from eight NLR genes that all reside in a defined NLR locus on the top of potato chromosome 5. At 5% and 7% mismatch rates, the RenSeq-derived SNPs corresponded to 58 SNPs in 15 NLRs and 75 SNPs in 18 NLRs (Fig. 3). With the exception of a single gene with informative SNPs on chromosome 8, all map to linkage group 5 (Tables S4, S5). By combining the NLRs linked to the resistance at the different mismatch rates, the putative resistance locus is estimated to span less than 12 MB between DM positions 4.2 MB to 16 MB on chromosome 5 (Table S5).

**Fig. 3.**
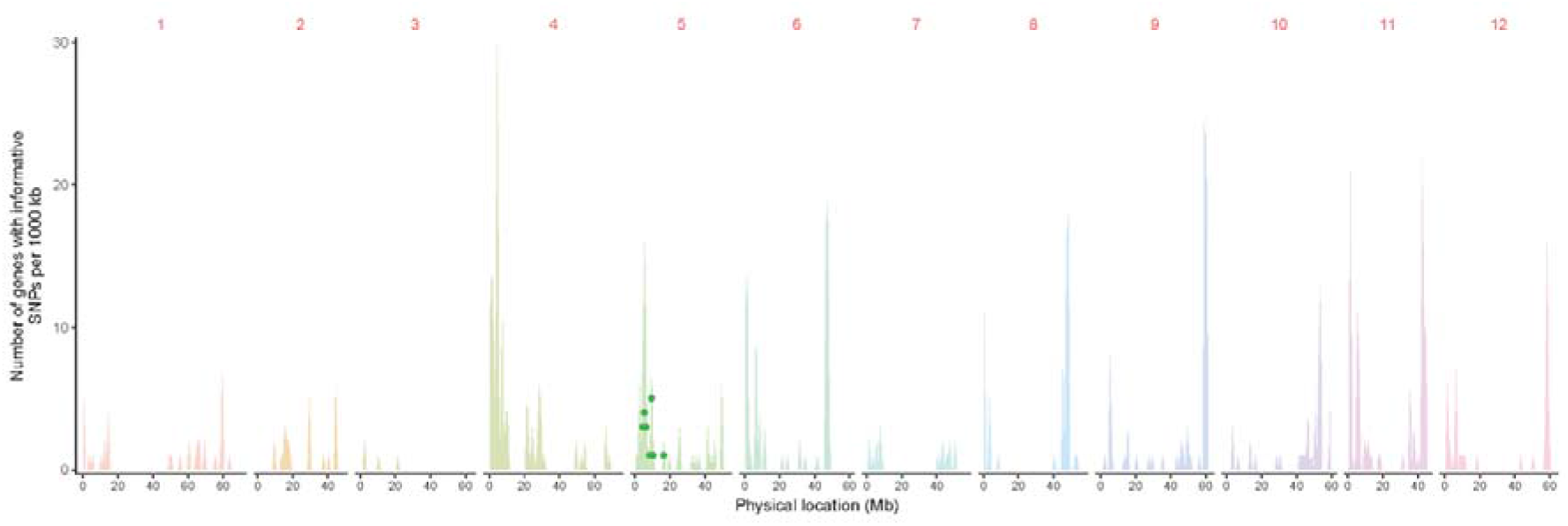
Graphical representation of NB-LRRs containing informative SNPs linked to *Rpi-blb4* at a 7 % mismatch rate. Chromosomes (1-12) are depicted on the x-axis and the number of genes with informative SNPs within 1Mb intervals are shown as dots on chromosome 5. Shaded in the background is the number of NLR genes that were assessed for linkage at each locus.

Similarly, the 10 GenSeq-derived SNPs at the 3% mismatch rate are derived from seven genes of which six correspond to potato linkage group 5 and a single SNP potentially corresponds to a gene on linkage group 10 (Table 2). However, at the 5% and 7% mismatch rates, all GenSeq-derived SNPs (57 SNPs in 28 genes and 66 SNPs in 32 genes), were unequivocally mapped to potato chromosome 5 (Tables S4, S5). Since *Rpi-blb1* has been located on chromosome 8 [30], *Rpi-blb2* on chromosome 6 [28] and *Rpi-blb3* on chromosome 4, these genetic mapping data support the hypothesis that *BLB*7650 contains a novel putative NLR-based resistance to *P. infestans*.

**Table 2.**
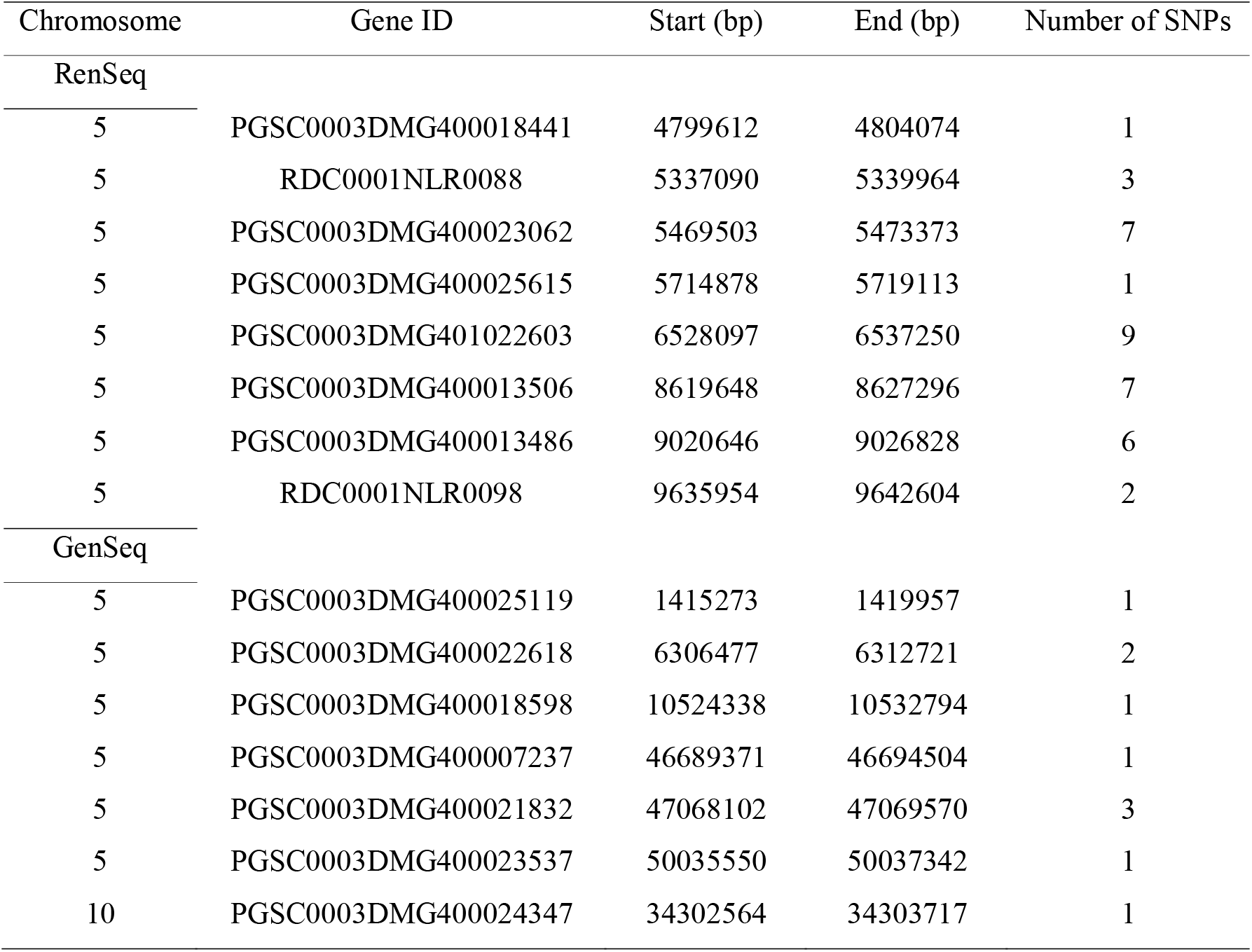
Positional mapping of SNPs from RenSeq and GenSeq analysis at a 3% mismatch rate using the reference potato genome *S. tuberosum* group Phureja clone DM1-3 516 R44. The corresponding gene names and their positions on the potato chromosomes are shown along with the number of informative SNPs per gene.

| Chromosome | Gene ID | Start (bp) | End (bp) | Number of SNPs |
| --- | --- | --- | --- | --- |
| RenSeq |  |  |  |  |
| 5 | PGSC0003DMG400018441 | 4799612 | 4804074 | 1 |
| 5 | RDC0001NLR0088 | 5337090 | 5339964 | 3 |
| 5 | PGSC0003DMG400023062 | 5469503 | 5473373 | 7 |
| 5 | PGSC0003DMG400025615 | 5714878 | 5719113 | 1 |
| 5 | PGSC0003DMG401022603 | 6528097 | 6537250 | 9 |
| 5 | PGSC0003DMG400013506 | 8619648 | 8627296 | 7 |
| 5 | PGSC0003DMG400013486 | 9020646 | 9026828 | 6 |
| 5 | RDC0001NLR0098 | 9635954 | 9642604 | 2 |
| GenSeq |  |  |  |  |
| 5 | PGSC0003DMG400025119 | 1415273 | 1419957 | 1 |
| 5 | PGSC0003DMG400022618 | 6306477 | 6312721 | 2 |
| 5 | PGSC0003DMG400018598 | 10524338 | 10532794 | 1 |
| 5 | PGSC0003DMG400007237 | 46689371 | 46694504 | 1 |
| 5 | PGSC0003DMG400021832 | 47068102 | 47069570 | 3 |
| 5 | PGSC0003DMG400023537 | 50035550 | 50037342 | 1 |
| 10 | PGSC0003DMG400024347 | 34302564 | 34303717 | 1 |

### 3.4. Graphical genotyping confirms the *Rpi-blb4* resistance on chromosome 5 and defines the locus to a 2.3 MB interval

To confirm the position of *Rpi-blb4* on Chromosome 5, selected RenSeq and GenSeq-derived SNPs linked to the resistance (Tables 2, S4, S5) were converted into allele-specific KASP markers. Initially, the progeny clones belonging to bulked resistant and bulked susceptible were assessed alongside the parents, BLB7650 and MCH3847. DNA isolated from fresh cuttings of the 23 resistant and 24 susceptible progeny clones (Table S3) were evaluated genotypically with allele-specific markers from Chromosome 5 NLR PGSC0003DMG401022603 at position 6533979 and the GenSeq-derived marker corresponding to PGSC0003DMG400019810 at position 15145865 (Table S6).

Consistent with the RenSeq and GenSeq analysis, plants in the resistant bulk contained mainly alleles from the resistant parent, BLB7650, and plants in the susceptible bulk alleles from MCH3847. This analysis further identified potential recombination events in the resistant bulk plants 3 and 5 and susceptible bulk plants 35, 36, and 52. One plant, number 74 from the susceptible bulk, however only contained markers associated with the resistant parent. This plant was re-phenotyped with the late blight isolate 2009-7654A and, in contrast to previous results (Table S3), displayed resistance, suggesting a mix-up of the plant during maintenance (Fig. 4A).

**Fig. 4.**
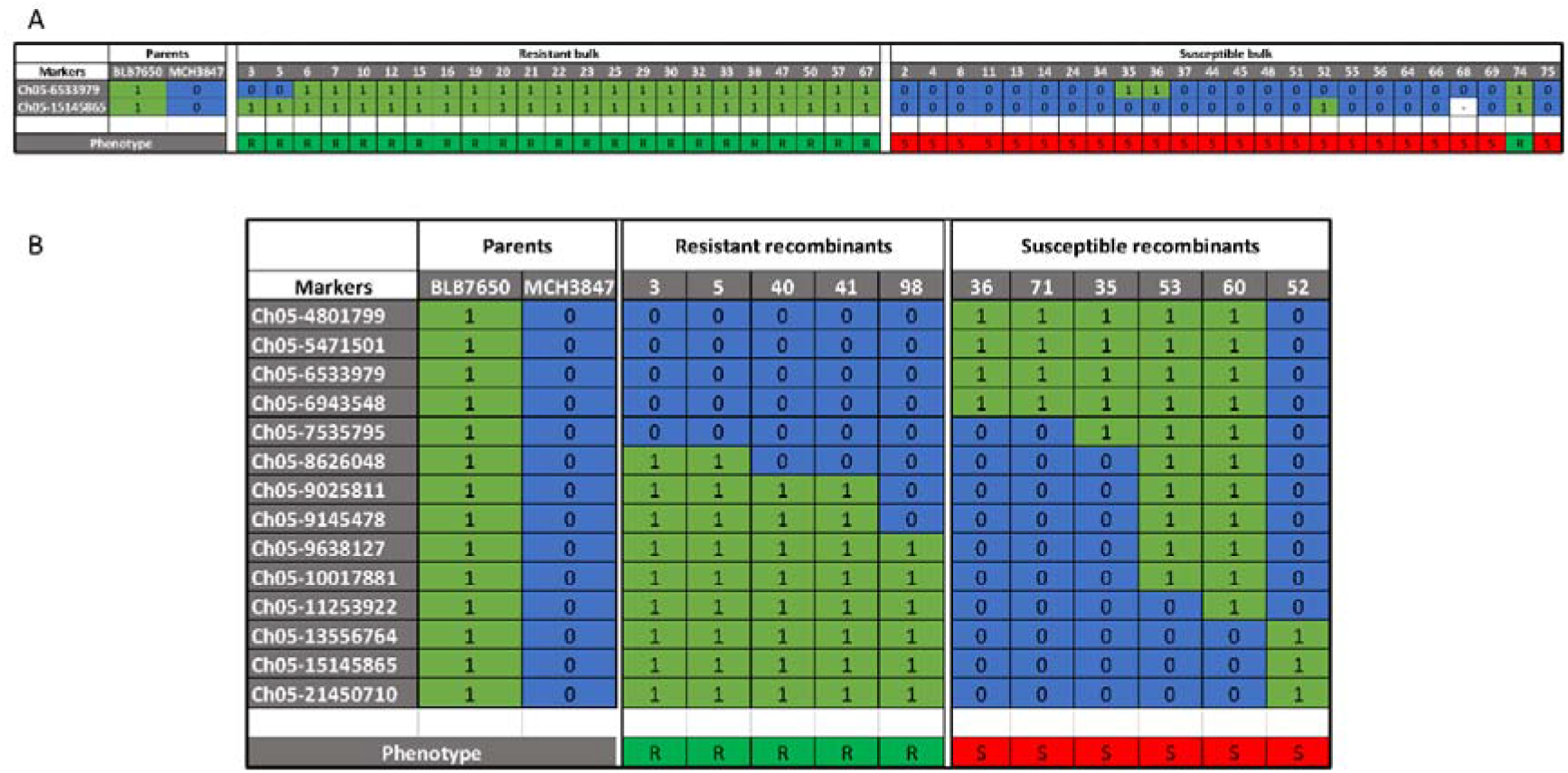
KASP-based mapping of *Rpi-blb4*. Initially, the RenSeq-based (DMG401022603) and GenSeq-based (DMG400019810) SNPs were assessed on the resistant and susceptible parents alongside the individual progeny clones that made up the bulks used in this study (A). Additional KASP markers were designed and utilised to narrow the *Rpi-blb4* interval in recombinant progeny clones (B).

To further identify recombinants, the same markers were applied to the entire progeny. This identified six further recombinants (resistant plants 40, 41, and 98; susceptible plants 53, 60, and 71) (Fig. 4B). The 11 recombinant plants from the population were characterised with 14 KASP markers corresponding to SNPs identified in the RenSeq and GenSeq analyses. The markers span potato chromosome 5 from DM genome position 4.8 MB to 21.5 MB (Table S6). The selected progeny clones were arranged according to their recombination breakpoints and their phenotype (Fig. 4B).

Using the gene order as established in DM, no double recombination events were identified suggesting that the gene order is largely conserved between DM and BLB7650. Further, this analysis identified a highly informative recombination event between plants 52 and 60 that positions the new resistance between the markers ch05-11253922 and ch05-13556764. These markers correspond to positions 11.25 MB and 13.56 MB and delimit the resistance interval to 2.3 MB.

## 4. Discussion

Wild potato species that are maintained in international germplasm collections represent an important natural resource and contain a wealth of important crop traits including disease resistance against unrelated pathogens [39]. The diploid wild species *S. bulbocastanum* is native to Central America and is widely known for its high resistance to *P. infestans* [40]. To date, only 5 *NLR* genes have been identified in *S. bulbocastanum* and mapped to chromosome 8 (*Rpi-blb1*; RB, *Rpi-bt1*), chromosome 6 (*Rpi-blb2*), and chromosome 4 (*Rpi-blb3*) [28-32]. In this study, we report on the identification and mapping of a novel resistance, *Rpi-blb4*, in *S. bulbocastanum* accession 7650 using RenSeq and GenSeq-based enrichment sequencing as applied previously to *Solanum verrucosum* [20]. The application of dRenSeq [22,24] proved highly informative in establishing the uniqueness of the observed resistance in BLB7650 (Fig. 1) and informed the commitment to a genetic study. The distribution of the resistance in the progeny clones closely followed a 1:1 ratio (Fig. 2) which is typically expected from a single dominant resistance gene or physically closely linked genes that are segregating in a diploid population.

*Rpi-blb4* maps to the top end of chromosome 5 and is, with *R1*, only the second major late blight gene that has been positioned to this linkage group [2,10]. Maturity, which is indirectly linked to late blight resistance has, however, also been mapped to chromosome 5 but in the form of a quantitative trait loci (QTL) [41]. Other functional disease resistances associated with chromosome 5 include, for example, those effective against potato cyst nematode (PCN) and viruses. Resistance against the PCN species *Globodera pallida* has been identified in the form of *H2* [42], *Gpa*^*V*^_*spl*_ *from S. sparsipilum* as well as *Gpa/GpaM1* from *S. spegazzinii* [43], and *Gpa5* from *S. vernei* [44,45]. Resistances on chromosome 5 effective against *G. rostochiensis* include *H1* [46], *GroVI* [47], and *Grp1* which is effective against both PCN species [48]. Further, *Rx2*, a gene effective against Potato Virus X has also been identified on chromosome 5 [49].

The next steps toward isolating the underpinning gene could include expanding the population to identify additional recombinants in the described interval between the two flanking markers ch05-11253922 and ch05-13556764 and to develop further markers located in this interval. In addition to such a fine-mapping approach, two further approaches could be used to identify likely candidates. First, with the advancement of phasing the genomes of heterozygous plants including tetraploid potatoes (e.g. Hoopes et al., 2022 [50]) combined with long-read sequencing technologies such as Oxford nanopore and/or PacBio, the two haplotypes of *S. bulbocastanum* 7650 could be established. The established markers that are linked to the resistance (Tables 2, S4, S5) could be used to select the haplotype that is in coupling whilst rejecting the haplotype in repulsion. Secondly, SMRT-AgRenSeq [25] could be used to assemble the NLRs of BLB7650 using workflows such as HISS [51] and genetically determine candidates using the existing bulks. The latter approach assumes that an NLR gene is responsible for the resistance, which is indeed likely considering that known potato genes effective against late blight typically belong to this gene family [2].

## Authors’ contributions

JL, MA and AK conducted the enrichment sequencing, the computational analyses to identify linked SNPs, and developed the KASP markers. BH conducted the late blight screening and KASP marker analysis. XW and IH conceived this work and secured funding. All authors have read and endorsed the manuscript.

## Ethics and Integrity Declarations

The authors declare that they have no competing interests.

## Acknowledgments

The authors acknowledge the Research/Scientific Computing teams at The James Hutton Institute and NIAB for providing computational resources and technical support for the “UK’s Crop Diversity Bioinformatics HPC” (BBSRC grant BB/S019669/1), use of which has contributed to the results reported within this paper.

## Sequence data availability

The RenSeq and GenSeq sequencing data derived from *S. bulbocastanum* accession 7650, *S. michoacanum* accession 3847, and the bulks are available at the European Nucleotide Archive (ENA) at EMBL-EBI as project PRJEB61409.

## Supplementary material

**Table S1**. Late blight scores for the resistant parent, *Solanum bulbocastanum* accession 7650, following inoculations with diverse *P. infestans* isolate using detached leaf assay. The disease severity was scored on a scale from 1 to 5 where 1 represents highly susceptible and 5 highly resistant.

**Table S2**. dRenSeq analysis showing NB-LRR coverage (%) in parents *Solanum bulbocastanum* accession 7650, *S. michoacanum* 3847, and resistant/susceptible progeny bulks. RenSeq-derived Illumina reads were mapped to the sequences of known NB-LRR genes with a highly sensitive mode.

**Table S3**. Replicated blight scores of 100 F1 progeny clones derived from a cross between *S. bulbocastanum* accession 7650 and *S. michoacanum accession* MCH3847. Replicated detach leaf assays were used to assess the resistance of progeny clones against the *P. infestans* isolate 2009-7654A. Blight severity is recorded according to the Malcolmson scale where 1 represents highly susceptible and 5 highly resistant. Average scores are calculated, and plants are classified as susceptible (red fields marked ‘S’ with scores between 1 and <2.5) intermediate (blue fields marked ‘/’with scores >2.5 and <3.5), and resistant (green fields marked ‘R’ with scores >3.5) Plants selected for bulked segregant analysis are highlighted for susceptible (red fields marked ‘BS’) and resistant (green fields marked ‘BR’) progeny clones.

**Table S4**. Positional mapping of SNPs from RenSeq and GenSeq analysis at a 5% mismatch rate compared to the potato reference genome (*S. tuberosum* group Phureja clone DM1-3 516 R44). The gene identification and position of NB-LRR containing filtered SNPs on the chromosome are shown along with the number of informative SNPs per gene.

**Table S5**. Positional mapping of SNPs from RenSeq and GenSeq analysis at a 7% mismatch rate compared to the potato genome (*S. tuberosum* group Phureja clone DM1-3 516 R44). The gene identification and position of NB-LRR containing filtered SNPs on the chromosome are shown along with the number of informative SNPs per gene.

**Table S6**. Description of the KASP markers used for the mapping of *Rpi-blb4*. Shown are the potato gene IDs, RenSeq or GenSeq origin, the corresponding mismatch rate used to identify the sequence polymorphism, the Assay ID (including the position in the DM1-3 516 R44 genome), the alleles, and corresponding primers used to assay the SNPs.

