## Supplementary Tables S1-S6 for "Identification and mapping of *Rpi-blb4* in diploid wild potato species *Solanum bulbocastanum*"

**Supplementary material**

**Table S1.** Blight scores for the resistant parent, *Solanum bulbocastanum* accession 7650, following inoculations with diverse *P. infestans* isolate using detached leaf assay. The disease severity was scored on a scale from 1 to 5 where 1 represents highly susceptible and 5 highly resistant.

| Isolate Origin | Isolate name | BLB7650 [replicates] | BLB7650 [average] |
| --- | --- | --- | --- |
| U.K. | 7804B [Other] | 5,5,5,5,5 | 5 |
| Netherlands | NL11564 | 5,4,5 | 4.6 |
| Netherlands | NL12206 | 5,5,5 | 5 |
| Netherlands | NL09096 | 2,2 | 2 |
| Netherlands | 08797 | 5,5,5,4 | 4.8 |
| Netherlands | 14538 | 5,3,5,5,4,5 | 4.5 |
| U.S.A. | US23 | 5,5,5 | 5 |
| U.K. | 1555C [13A2] | 5,5,5,5,5,5,5,4,4 | 4.8 |
| Netherlands | 14534 | 5,4,4 | 4.3 |
| U.K. | W3872A [13A2] | 5,5 | 5 |
| U.K. | W8814B [13A2] | 5,5 | 5 |
| U.K. | W4425B [13A2] | 4,4 | 4 |
| U.K. | W4615A [13A2] | 4,5,4,4 | 4.3 |
| U.K. | W10421A [36A2] | 5,4,5,4,4,5 | 4.5 |
| U.K. | W9928C [36A2] | 5,5,4,5,5 | 4.8 |

**Table S2.** dRenSeq analysis showing NB-LRR coverage (%) in parents *Solanum bulbocastanum* accession 7650, *S. michoacanum* 3847 and resistant/susceptible progeny bulks. RenSeq-derived Illumina reads were mapped to the sequences of known NB-LRR genes with a highly sensitive mode.

| Gene Names | BLB7650 | MCH3847 | Bulk Resistant | Bulk Susceptible |
| --- | --- | --- | --- | --- |
| *Rpi-abpt* | 55.0433 | 16.1939 | 45.6659 | 56.383 |
| *Rpi-amr1* | 0 | 0 | 0 | 0 |
| *Rpi-amr3* | 0 | 0 | 0 | 0 |
| *Rpi-ber* | 0 | 0 | 0 | 0 |
| *Rpi-ber1.1* | 0 | 0 | 0 | 0 |
| *Rpi-ber1.2* | 0 | 0 | 0 | 0 |
| *Rpi-blb1* | 0 | 0 | 0 | 0 |
| *Rpi-blb2* | 16.9923 | 6.58098 | 11.7995 | 13.1362 |
| *Rpi-blb3* | 100 | 8.13679 | 100 | 100 |
| *Rpi-bt1* | 4.91415 | 0 | 0 | 0 |
| *Rpi-chc1.1* | 0 | 0 | 0 | 0 |
| *Rpi-chc1.2* | 0 | 0 | 0 | 0 |
| *Rpi-hcb1.1* | 0 | 0 | 0 | 0 |
| *Rpi-hcb1.2* | 0 | 0 | 0 | 0 |
| *Rpi-mcq1.1* | 0 | 0 | 0 | 0 |
| *Rpi-pta1* | 0 | 0 | 0 | 0 |
| *Rpi-R1* | 0 | 0 | 0 | 0 |
| *Rpi-R2* | 32.3877 | 8.15603 | 26.3987 | 27.7384 |
| *Rpi-R2-like* | 69.1431 | 16.1557 | 60.9277 | 69.9686 |
| *Rpi-R3a* | 10.0026 | 10.4703 | 6.00156 | 10.4962 |
| *Rpi-R3b* | 30.5036 | 52.8557 | 47.8972 | 56.9315 |
| *Rpi-R8* | 3.18352 | 0 | 0 | 0 |
| *Rpi-R9a* | 0 | 0 | 0 | 0 |
| *Rpi-sto1* | 0 | 0 | 0 | 0 |
| *Rpi-tar1* | 0 | 0 | 0 | 0 |
| *Rpi-vnt1.1* | 0 | 0 | 0 | 0 |
| *Rpi-vnt1.3* | 0 | 0 | 0 | 0 |

**Table S3.** Replicated blight scores of 100 F1 progeny clones derived from a cross between *S. bulbocastanum* accession 7650 and *S. michoacanum accession* MCH3847. Replicated detach leaf assays were used to assess the resistance of progeny clones against the *P. infestans* isolate 2009-7654A. Blight severity is recorded according to the Malcolmson scale where 1 represents highly susceptible and 5 highly resistant. Average scores are calculated and plants are classified as susceptible (red fields marked ‘S’ with scores between 1 and <2.5) intermediate (blue fields marked ‘/’with scores >2.5 and <3.5) and resistant (green fields marked ‘R’ with scores >3.5) Plants selected for bulked segregant analysis are highlighted for susceptible (red fields marked ‘BS’) and resistant (green fields marked ‘BR’) progeny clones.

| Plant Number | Detached Leaves scores | | | | | | | | | | Average | Resistant | Susceptible | Intermediate | TOP 23 RES | TOP 24 SUS |
| --- | --- | --- | --- | --- | --- | --- | --- | --- | --- | --- | --- | --- | --- | --- | --- | --- |
|  | 1 | 2 | 3 | 4 | 5 | 6 | 7 | 8 | 9 | 10 |  |  |  |  |  |  |
| 1 | 1 | 2 | 3 | 2 | 5 | 5 | 1 | 1 | 2 | 1 | 2.3 |  | S |  |  |  |
| 2 | 2 | 2 | 3 | 1 | 1 | 1 |  |  |  |  | 1.666667 |  | S |  |  | BS |
| 3 | 4 | 5 | 5 | 4 | 5 |  |  |  |  |  | 4.6 | R |  |  | BR |  |
| 4 | 1 | 1 | 3 | 3 | 1 | 2 | 1 | 1 |  |  | 1.625 |  | S |  |  | BS |
| 5 | 5 | 4 | 5 | 4 | 5 |  |  |  |  |  | 4.6 | R |  |  | BR |  |
| 6 | 5 | 5 | 4 | 5 |  |  |  |  |  |  | 4.75 | R |  |  | BR |  |
| 7 | 5 | 4 | 4 | 4 | 5 |  |  |  |  |  | 4.4 | R |  |  | BR |  |
| 8 | 3 | 2 | 1 | 2 | 1 | 1 | 3 | 1 |  |  | 1.75 |  | S |  |  | BS |
| 9 | 2 | 1 | 5 | 5 | 5 | 1 | 3 | 1 | 3 |  | 2.888889 |  |  | / |  |  |
| 10 | 5 | 5 | 5 | 5 |  |  |  |  |  |  | 5 | R |  |  | BR |  |
| 11 | 2 | 1 | 3 | 1 | 2 | 2 | 3 | 1 |  |  | 1.875 |  | S |  |  | BS |
| 12 | 5 | 5 | 5 | 5 | 4 |  |  |  |  |  | 4.8 | R |  |  | BR |  |
| 13 | 2 | 1 | 1 | 2 | 3 |  |  |  |  |  | 1.8 |  | S |  |  | BS |
| 14 | 2 | 1 | 1 | 1 | 2 |  |  |  |  |  | 1.4 |  | S |  |  | BS |
| 15 | 5 | 5 | 5 |  |  |  |  |  |  |  | 5 | R |  |  | BR |  |
| 16 | 5 | 5 | 4 |  |  |  |  |  |  |  | 4.666667 | R |  |  | BR |  |
| 17 | 1 | 3 | 1 | 5 | 5 | 3 | 1 |  |  |  | 2.714286 |  |  | / |  |  |
| 18 | 2 | 2 | 3 | 4 | 4 | 5 | 2 | 1 |  |  | 2.875 |  |  | / |  |  |
| 19 | 5 | 5 | 5 |  |  |  |  |  |  |  | 5 | R |  |  | BR |  |
| 20 | 5 | 5 | 5 |  |  |  |  |  |  |  | 5 | R |  |  | BR |  |
| 21 | 5 | 5 | 5 | 4 |  |  |  |  |  |  | 4.75 | R |  |  | BR |  |
| 22 | 5 | 5 | 5 | 5 |  |  |  |  |  |  | 5 | R |  |  | BR |  |
| 23 | 4 | 4 | 4 | 4 |  |  |  |  |  |  | 4 | R |  |  | BR |  |
| 24 | 2 | 1 | 1 | 1 | 1 |  |  |  |  |  | 1.2 |  | S |  |  | BS |
| 25 | 5 | 5 | 4 |  |  |  |  |  |  |  | 4.666667 | R |  |  | BR |  |
| 26 | 3 | 2 | 3 | 3 | 4 | 5 |  |  |  |  | 3.333333 |  |  | / |  |  |
| 27 | 4 | 2 | 2 | 3 | 2 | 2 |  |  |  |  | 2.5 |  |  | / |  |  |
| 28 | 4 | 2 | 4 | 5 |  |  |  |  |  |  | 3.75 | R |  |  |  |  |
| 29 | 4 | 4 | 5 |  |  |  |  |  |  |  | 4.333333 | R |  |  | BR |  |
| 30 | 3 | 4 | 4 | 4 | 5 | 5 |  |  |  |  | 4.166667 | R |  |  | BR |  |
| 31 | 3 | 4 | 4 | 4 |  |  |  |  |  |  | 3.75 | R |  |  |  |  |
| 32 | 3 | 4 | 5 | 3 | 5 | 5 |  |  |  |  | 4.166667 | R |  |  | BR |  |
| 33 | 3 | 5 | 5 | 4 | 5 | 5 |  |  |  |  | 4.5 | R |  |  | BR |  |
| 34 | 1 | 1 | 1 | 1 |  |  |  |  |  |  | 1 |  | S |  |  | BS |
| 35 | 1 | 1 | 2 | 1 | 2 |  |  |  |  |  | 1.4 |  | S |  |  | BS |
| 36 | 1 | 1 | 1 | 1 | 2 |  |  |  |  |  | 1.2 |  | S |  |  | BS |
| 37 | 1 | 1 | 3 | 1 |  |  |  |  |  |  | 1.5 |  | S |  |  | BS |
| 38 | 5 | 5 | 4 |  |  |  |  |  |  |  | 4.666667 | R |  |  | BR |  |
| 39 | 3 | 4 | 4 | 4 |  |  |  |  |  |  | 3.75 | R |  |  |  |  |
| 40 | 3 | 4 | 3 | 4 | 5 | 4 | 3 | 3 |  |  | 3.625 | R |  |  |  |  |
| 41 | 2 | 5 | 4 | 4 | 5 | 4 | 5 |  |  |  | 4.142857 | R |  |  |  |  |
| 42 | 1 | 1 | 3 |  |  |  |  |  |  |  | 1.666667 |  | S |  |  |  |
| 43 | 3 | 4 | 5 | 4 |  |  |  |  |  |  | 4 | R |  |  |  |  |
| 44 | 2 | 1 | 2 | 1 |  |  |  |  |  |  | 1.5 |  | S |  |  | BS |
| 45 | 1 | 1 | 1 | 1 |  |  |  |  |  |  | 1 |  | S |  |  | BS |
| 46 | 1 | 2 | 4 | 4 | 3 | 2 | 4 | 3 | 2 |  | 2.777778 |  |  | / |  |  |
| 47 | 5 | 5 | 4 |  |  |  |  |  |  |  | 4.666667 | R |  |  | BR |  |
| 48 | 1 | 1 | 1 |  |  |  |  |  |  |  | 1 |  | S |  |  | BS |
| 49 | 1 | 1 | 3 |  |  |  |  |  |  |  | 1.666667 |  | S |  |  |  |
| 50 | 5 | 5 | 4 |  |  |  |  |  |  |  | 4.666667 | R |  |  | BR |  |
| 51 | 1 | 1 | 1 |  |  |  |  |  |  |  | 1 |  | S |  |  | BS |
| 52 | 1 | 1 | 1 |  |  |  |  |  |  |  | 1 |  | S |  |  | BS |
| 53 | 3 | 2 | 4 | 4 | 4 | 5 | 2 |  |  |  | 3.428571 |  |  | / |  |  |
| 54 | 1 | 1 | 3 | 3 |  |  |  |  |  |  | 2 |  | S |  |  |  |
| 55 | 1 | 1 | 1 | 1 |  |  |  |  |  |  | 1 |  | S |  |  | BS |
| 56 | 1 | 1 | 1 |  |  |  |  |  |  |  | 1 |  | S |  |  | BS |
| 57 | 5 | 4 | 4 |  |  |  |  |  |  |  | 4.333333 | R |  |  | BR |  |
| 58 | 4 | 3 | 3 | 4 | 5 | 5 |  |  |  |  | 4 | R |  |  |  |  |
| 59 | 5 | 3 | 2 | 3 | 3 | 2 | 3 |  |  |  | 3 |  |  | / |  |  |
| 60 | 3 | 1 | 2 | 3 | 3 | 3 | 1 | 2 |  |  | 2.25 |  | S |  |  |  |
| 61 | 3 | 3 | 5 | 5 | 4 | 5 |  |  |  |  | 4.166667 | R |  |  |  |  |
| 62 | 1 | 3 | 3 |  |  |  |  |  |  |  | 2.333333 |  | S |  |  |  |
| 63 | 2 | 1 | 3 | 3 | 4 | 5 | 4 | 4 |  |  | 3.25 |  |  | / |  |  |
| 64 | 1 | 1 | 1 |  |  |  |  |  |  |  | 1 |  | S |  |  | BS |
| 65 | 2 | 2 | 3 | 3 | 2 | 2 |  |  |  |  | 2.333333 |  | S |  |  |  |
| 66 | 1 | 1 | 1 |  |  |  |  |  |  |  | 1 |  | S |  |  | BS |
| 67 | 4 | 4 | 5 |  |  |  |  |  |  |  | 4.333333 | R |  |  | BR |  |
| 68 | 1 | 1 | 1 |  |  |  |  |  |  |  | 1 |  | S |  |  | BS |
| 69 | 1 | 1 | 1 |  |  |  |  |  |  |  | 1 |  | S |  |  | BS |
| 70 | 4 | 1 | 4 | 4 |  |  |  |  |  |  | 3.25 |  |  | / |  |  |
| 71 | 1 | 1 | 3 |  |  |  |  |  |  |  | 1.666667 |  | S |  |  |  |
| 72 | 4 | 2 | 4 | 4 | 4 |  |  |  |  |  | 3.6 | R |  |  |  |  |
| 73 | 3 | 2 | 2 | 4 | 3 | 5 |  |  |  |  | 3.166667 |  |  | / |  |  |
| 74 | 2 | 2 | 1 | 1 | 2 | 4 |  |  |  |  | 2 |  | S |  |  | BS |
| 75 | 1 | 1 | 1 | 1 |  |  |  |  |  |  | 1 |  | S |  |  | BS |
| 76 | 1 | 1 |  |  |  |  |  |  |  |  | 1 |  | S |  |  |  |
| 77 | 1 | 4 |  |  |  |  |  |  |  |  | 2.5 |  |  | / |  |  |
| 78 | 1 |  |  |  |  |  |  |  |  |  | 1 |  | S |  |  |  |
| 79 | 1 |  |  |  |  |  |  |  |  |  | 1 |  | S |  |  |  |
| 80 | 5 | 3 |  |  |  |  |  |  |  |  | 4 | R |  |  |  |  |
| 81 | 5 | 2 | 2 |  |  |  |  |  |  |  | 3 |  |  | / |  |  |
| 82 | 5 | 5 | 5 |  |  |  |  |  |  |  | 5 | R |  |  |  |  |
| 83 | 5 | 3 | 2 |  |  |  |  |  |  |  | 3.333333 |  |  | / |  |  |
| 84 | 4 | 3 | 2 |  |  |  |  |  |  |  | 3 |  |  | / |  |  |
| 85 | 5 | 5 |  |  |  |  |  |  |  |  | 5 | R |  |  |  |  |
| 86 | 5 | 5 |  |  |  |  |  |  |  |  | 5 | R |  |  |  |  |
| 87 | 5 | 5 |  |  |  |  |  |  |  |  | 5 | R |  |  |  |  |
| 88 | 3 | 2 | 3 | 5 |  |  |  |  |  |  | 3.25 |  |  | / |  |  |
| 89 | 5 | 5 | 5 |  |  |  |  |  |  |  | 5 | R |  |  |  |  |
| 90 | 5 | 5 | 5 |  |  |  |  |  |  |  | 5 | R |  |  |  |  |
| 91 | 5 | 1 | 1 | 3 |  |  |  |  |  |  | 2.5 |  |  | / |  |  |
| 92 | 5 | 2 | 3 | 5 |  |  |  |  |  |  | 3.75 | R |  |  |  |  |
| 93 | 5 | 4 | 4 |  |  |  |  |  |  |  | 4.333333 | R |  |  |  |  |
| 94 | 5 | 4 | 4 | 2 | 1 | 1 |  |  |  |  | 2.833333 |  |  | / |  |  |
| 95 | 5 | 4 | 4 | 4 | 1 | 1 |  |  |  |  | 3.166667 |  |  | / |  |  |
| 96 | 5 | 5 | 5 |  |  |  |  |  |  |  | 5 | R |  |  |  |  |
| 97 | 5 | 5 |  |  |  |  |  |  |  |  | 5 | R |  |  |  |  |
| 98 | 5 | 3 | 4 | 3 | 4 |  |  |  |  |  | 3.8 | R |  |  |  |  |
| 99 | 1 | 1 | 1 | 2 | 1 | 1 |  |  |  |  | 1.166667 |  | S |  |  |  |
| 100 | 5 | 5 | 5 |  |  |  |  |  |  |  | 5 | R |  |  |  |  |

**Table S4.** Positional mapping of SNPs from RenSeq and GenSeq analysis at 5% mismatch compared to the potato reference genome (*S. tuberosum* group Phureja clone DM1-3 516 R44). The gene identification and position of NB-LRR containing filtered SNPs on the chromosome are shown along with the number of informative SNPs per gene.

| Chromosome | Gene ID | Start (bp) | End (bp) | Number of SNPs |
| --- | --- | --- | --- | --- |
| RenSeq |  |  |  |  |
| 5 | RDC0001NLR0077 | 4232244 | 4235519 | 1 |
| 5 | PGSC0003DMG400018441 | 4799612 | 4804074 | 2 |
| 5 | RDC0001NLR0088 | 5337090 | 5339964 | 1 |
| 5 | PGSC0003DMG400023062 | 5469503 | 5473373 | 10 |
| 5 | PGSC0003DMG400025615 | 5714878 | 5719113 | 2 |
| 5 | PGSC0003DMG400025611 | 5723483 | 5731577 | 9 |
| 5 | PGSC0003DMG401022603 | 6528097 | 6537250 | 10 |
| 5 | RDC0001NLR0093 | 6938661 | 6944482 | 4 |
| 5 | PGSC0003DMG400013506 | 8619648 | 8627296 | 8 |
| 5 | PGSC0003DMG400013486 | 9020646 | 9026828 | 1 |
| 5 | PGSC0003DMG400013490 | 9139914 | 9148796 | 2 |
| 5 | RDC0001NLR0094 | 9151974 | 9154278 | 2 |
| 5 | PGSC0003DMG401013522 | 9287263 | 9291478 | 1 |
| 5 | RDC0001NLR0098 | 9635954 | 9642604 | 3 |
| 8 | PGSC0003DMG400012466 | 773509 | 780723 | 2 |
| GenSeq |  |  |  |  |
| 5 | PGSC0003DMG400025119 | 1415273 | 1419957 | 1 |
| 5 | PGSC0003DMG400025121 | 1437168 | 1441274 | 1 |
| 5 | PGSC0003DMG400018405 | 4484319 | 4492247 | 1 |
| 5 | PGSC0003DMG400018411 | 4734615 | 4737869 | 1 |
| 5 | PGSC0003DMG400003384 | 5359366 | 5362349 | 1 |
| 5 | PGSC0003DMG400022618 | 6306477 | 6312721 | 9 |
| 5 | PGSC0003DMG400022610 | 6729414 | 6734055 | 1 |
| 5 | PGSC0003DMG400022615 | 6769503 | 6774474 | 1 |
| 5 | PGSC0003DMG400004123 | 7531834 | 7537250 | 1 |
| 5 | PGSC0003DMG400004129 | 7620910 | 7623275 | 1 |
| 5 | PGSC0003DMG400030998 | 8383814 | 8387263 | 1 |
| 5 | PGSC0003DMG400030985 | 8404032 | 8405976 | 7 |
| 5 | PGSC0003DMG400037297 | 10017565 | 10018209 | 1 |
| 5 | PGSC0003DMG400018621 | 10063175 | 10068988 | 1 |
| 5 | PGSC0003DMG400018598 | 10524338 | 10532794 | 1 |
| 5 | PGSC0003DMG400011728 | 10597983 | 10604575 | 1 |
| 5 | PGSC0003DMG400011727 | 10614999 | 10623108 | 2 |
| 5 | PGSC0003DMG400010739 | 11252622 | 11256056 | 3 |
| 5 | PGSC0003DMG400015053 | 13553252 | 13557860 | 8 |
| 5 | PGSC0003DMG400019810 | 15144696 | 15151519 | 1 |
| 5 | PGSC0003DMG400019812 | 15340984 | 15341956 | 2 |
| 5 | PGSC0003DMG400041031 | 21448468 | 21451136 | 2 |
| 5 | PGSC0003DMG400007038 | 44913810 | 44918918 | 1 |
| 5 | PGSC0003DMG400006987 | 45667733 | 45676099 | 1 |
| 5 | PGSC0003DMG400007222 | 46583390 | 46590258 | 1 |
| 5 | PGSC0003DMG400040438 | 46658289 | 46659272 | 1 |
| 5 | PGSC0003DMG400007237 | 46689371 | 46694504 | 1 |
| 5 | PGSC0003DMG400021832 | 47068102 | 47069570 | 4 |

**Table S5.** Positional mapping of SNPs from RenSeq and GenSeq analysis at 7% mismatch compared to the potato genome (*S. tuberosum* group Phureja clone DM1-3 516 R44). The gene identification and position of NB-LRR containing filtered SNPs on the chromosome is shown along with the number of informative SNPs per gene.

| Chromosome | Gene ID | Start (bp) | End (bp) | Number of SNPs |
| --- | --- | --- | --- | --- |
| RenSeq |  |  |  |  |
| 5 | PGSC0003DMG400030497 | 4227604 | 4230353 | 1 |
| 5 | RDC0001NLR0077 | 4232244 | 4235519 | 1 |
| 5 | PGSC0003DMG400018441 | 4799612 | 4804074 | 2 |
| 5 | RDC0001NLR0088 | 5337090 | 5339964 | 1 |
| 5 | PGSC0003DMG400023062 | 5469503 | 5473373 | 10 |
| 5 | PGSC0003DMG400025615 | 5714878 | 5719113 | 2 |
| 5 | PGSC0003DMG400025611 | 5723483 | 5731577 | 8 |
| 5 | RDC0001NLR0090 | 6506321 | 6508868 | 1 |
| 5 | PGSC0003DMG401022603 | 6528097 | 6537250 | 10 |
| 5 | RDC0001NLR0093 | 6938661 | 6944482 | 6 |
| 5 | PGSC0003DMG400013506 | 8619648 | 8627296 | 18 |
| 5 | PGSC0003DMG400013486 | 9020646 | 9026828 | 1 |
| 5 | PGSC0003DMG400013490 | 9139914 | 9148796 | 1 |
| 5 | RDC0001NLR0094 | 9151974 | 9154278 | 3 |
| 5 | PGSC0003DMG401013522 | 9287263 | 9291478 | 3 |
| 5 | RDC0001NLR0098 | 9635954 | 9642604 | 3 |
| 5 | PGSC0003DMG400018619 | 10091268 | 10095994 | 2 |
| 5 | PGSC0003DMG400002357 | 16044827 | 16048908 | 2 |
| GenSeq |  |  |  |  |
| 5 | PGSC0003DMG400025119 | 1415273 | 1419957 | 1 |
| 5 | PGSC0003DMG400025121 | 1437168 | 1441274 | 1 |
| 5 | PGSC0003DMG400018405 | 4484319 | 4492247 | 1 |
| 5 | PGSC0003DMG400037297 | 10017565 | 10018209 | 1 |
| 5 | PGSC0003DMG400018621 | 10063175 | 10068988 | 1 |
| 5 | PGSC0003DMG400018598 | 10524338 | 10532794 | 1 |
| 5 | PGSC0003DMG400011728 | 10597983 | 10604575 | 1 |
| 5 | PGSC0003DMG400011727 | 10614999 | 10623108 | 2 |
| 5 | PGSC0003DMG400010739 | 11252622 | 11256056 | 3 |
| 5 | PGSC0003DMG400015053 | 13553252 | 13557860 | 5 |
| 5 | PGSC0003DMG400019810 | 15144696 | 15151519 | 1 |
| 5 | PGSC0003DMG400019812 | 15340984 | 15341956 | 3 |
| 5 | PGSC0003DMG400041031 | 21448468 | 21451136 | 2 |
| 5 | PGSC0003DMG400018834 | 33928253 | 33930071 | 1 |
| 5 | PGSC0003DMG400007038 | 44913810 | 44918918 | 1 |
| 5 | PGSC0003DMG400006987 | 45667733 | 45676099 | 1 |
| 5 | PGSC0003DMG400007222 | 46583390 | 46590258 | 3 |
| 5 | PGSC0003DMG400040438 | 46658289 | 46659272 | 2 |
| 5 | PGSC0003DMG400007237 | 46689371 | 46694504 | 1 |
| 5 | PGSC0003DMG400021832 | 47068102 | 47069570 | 3 |
| 5 | PGSC0003DMG400018411 | 4734615 | 4737869 | 1 |
| 5 | PGSC0003DMG400003384 | 5359366 | 5362349 | 1 |
| 5 | PGSC0003DMG400025609 | 5679823 | 5684334 | 1 |
| 5 | PGSC0003DMG401017626 | 5918681 | 5923232 | 2 |
| 5 | PGSC0003DMG400022618 | 6306477 | 6312721 | 11 |
| 5 | PGSC0003DMG400022610 | 6729414 | 6734055 | 1 |
| 5 | PGSC0003DMG400004123 | 7531834 | 7537250 | 1 |
| 5 | PGSC0003DMG400004129 | 7620910 | 7623275 | 4 |
| 5 | PGSC0003DMG400030998 | 8383814 | 8387263 | 1 |
| 5 | PGSC0003DMG400030985 | 8404032 | 8405976 | 5 |
| 5 | PGSC0003DMG400013474 | 8659817 | 8660600 | 2 |
| 5 | PGSC0003DMG400013475 | 8663454 | 8668307 | 1 |

**Table S6.** Description of the KASP markers used for the mapping of *Rpi-blb4*. Shown are the potato gene IDs, RenSeq or GenSeq origin, the corresponding mismatch rate used to identify the sequence polymorphism, the Assay ID (including the position in the DM1-3 516 R44 genome), the alleles, and corresponding primers used to assay the SNPs.

| Gene | Type | Assays ID | FAM Allele | | HEX Allele | | Primer Seq Allele X | Primer Seq Allele Y | Primer Seq common |
| --- | --- | --- | --- | --- | --- | --- | --- | --- | --- |
| PGSC0003DMG400018441 | RenSeq 5% | ch05-4801799 | | A | | G | CCTCTCCATCCCAGCGAGTATA | CTCTCCATCCCAGCGAGTATG | TGTGAAGGGTTGCCTCTTTCAATTGTTTT |
| PGSC0003DMG400023062 | RenSeq 5% | ch05-5471501 | | C | | T | GATAATTCTACTTCCATTATTGGAATCATG | GGATAATTCTACTTCCATTATTGGAATCATA | GGAAGCTAGTGTTTGGGATGATTTAAGAT |
| PGSC0003DMG401022603 | RenSeq 3% | ch05-6533979 | | A | | G | CATGCATTTAGAAGACCAGACCCA | ATGCATTTAGAAGACCAGACCCG | CCAGTAATTGAGACAATTTGCTTGGATAAA |
| RDC0001NLR0093 | RenSeq 5% | ch05-6943548 | | C | | T | CCGCATCAATGTTTTGAGCAGATAC | ACCGCATCAATGTTTTGAGCAGATAT | GGTGTAYCTTAGAAATAAAAAACCTAAGAA |
| PGSC0003DMG400004123 | GenSeq 5% | ch05-7535795 | | A | | T | GAAGTCCCTATGAATTATAGGAATTCCA | GAAGTCCCTATGAATTATAGGAATTCCT | AAGAGGACTTGCATATCTGCATTCAAGTT |
| PGSC0003DMG400013506 | RenSeq 3% | ch05-8626048 | | G | | A | GACTGGGATCTATTGATATCATTTTAGAC | GACTGGGATCTATTGATATCATTTTAGAT | AGAAAGTGAGTCTGAATAGAGGMTTAGATA |
| PGSC0003DMG400013486 | RenSeq 5% | ch05-9025811 | | C | | G | CGTTCTTTGCTATTCAATGCCAGC | CGTTCTTTGCTATTCAATGCCAGG | TTAATGATGAAGGAGATATCAYGTGCCATT |
| PGSC0003DMG400013490 | RenSeq 5% | ch05-9145478 | | A | | T | CTGTCTCTTTCCCATCAGGGATA | CTGTCTCTTTCCCATCAGGGATT | AGCMTGTAACACCAAGAAGGTGCAT |
| RDC0001NLR0098 | RenSeq 3% | ch05-9638127 | | T | | C | CATATTCCCAATATGGATGAGATCTTGAT | ATTCCCAATATGGATGAGATCTTGAC | CATCCTTCCGWAGCAGATTGACAAGAA |
| PGSC0003DMG400037297 | GenSeq 5% | ch05-10017881 | | G | | A | GGATACTTCTACGATACATTGTACAC | CGGATACTTCTACGATACATTGTACAT | AGGAAATGTATCGCCRGGCTGCAA |
| PGSC0003DMG400010739 | GenSeq 5% | ch05-11253922 | | T | | C | TTTAGAAAGTGTGTTTCATTGTCTTGA | CTTTTAGAAAGTGTGTTTCATTGTCTTGG | GTAATTTGGTATATRAAGTCACTTAGTC |
| PGSC0003DMG400015053 | GenSeq 5% | ch05-13556764 | | A | | T | GTGTATGTTTGATGACCAAGTAKTCTTATA | GTGTATGTTTGATGACCAAGTAKTCTTATT | GCCAATGCCAAATTTAARCAACC |
| PGSC0003DMG400019810 | GenSeq 5% | ch05-15145865 | | C | | T | ATCTACATCACATCTCCTTGGGATG | AAATCTACATCACATCTCCTTGGGATA | CAACGTTCATGCAAGATCCAGGGAA |
| PGSC0003DMG400041031 | GenSeq 5% | ch05-21450710 | | T | | A | ATGGTATAATATTATAGGTTTGATAGTAAAGGT | ATGGTATAATATTATAGGTTTGATAGTAAAGGA | CCATCTATAGGAGCTATGGAGARATTTATA |
